# Analyzing ecological networks of species interactions

**DOI:** 10.1101/112540

**Authors:** Eva Delmas, Mathilde Besson, Marie-Hélène Brice, Laura A. Burkle, Giulio V. Dalla Riva, Marie-Josée Fortin, Dominique Gravel, Paulo R Guimarães, David Hembry, Erica Newman, Jens M. Olesen, Mathias M. Pires, Justin D. Yeakel, Timothée Poisot

## Abstract

Networks provide one of the best representations for ecological communities, composed of many species with sometimes complex connections between them. Yet the methodological literature allowing one to analyze and extract meaning from ecological networks is dense, fragmented, and unwelcoming. We provide a general overview to the field of using networks in community ecology, outlining both the intent of the different measures, their assumptions, and the contexts in which they can be used. When methodologically justified, we suggest good practices to use in the analysis of ecological networks. We anchor this synopsis with examples from empirical studies, and conclude by highlighting what identified as needed future developments in the field.

---

Abu ‘Uthman ‘Amr ibn Bahr was perhaps the first scientist to provide, as early as in the eighth century, a description of a food chain (Egerton 2002). About a thousand years later, Camerano (1880) introduced the idea that the diversity of animal forms, and therefore the biological diversity itself, can only be explained when framed in the context of inter-relationships between species. Seminal work by Patten (1978) or Ulanowicz (1980) suggested that the structure of networks can approximate information theoretical constraints on community assembly, and help generate interest in network science application for ecology. “Network-thinking” now permeates ecology and evolution (Proulx et al. 2005), and is one of the fastest growing ecological disciplines (Borrett et al. 2014), accounting for 5% of all published papers in 2012. Network-based approaches are gaining momentum as one of the most helpful tools for the analysis of community structure (Poisot et al. 2016d), because they offer the opportunity to investigate, within a common formal mathematical framework, questions ranging from the species-level to the community-level (Poisot et al. 2016d). Applying network approaches to a variety of ecological systems, for example hosts and parasites (Poulin 2010), or bacteria and phage (Weitz et al. 2013), yields new methodological and biological insights, such as the observation that networks tend to be locally nested but regionally modular (Flores et al. 2013), which suggests that different ecological and evolutionary regimes are involved at different scales. Despite this long-standing interest, the application of measures grounded in network science is still a relatively young field (in part because the computational power to perform some of these analysis was largely unavailable in the early days of the field). This comes with challenges to tackle. First, there is a pressing need for additional methodological developments, both to ensure that our quantitative analysis of networks is correct, and that it adequately captures the ecological realities that are, ultimately, of interest. Second, we need to better understand the limitations and domain of application of current methods. Yet, there is a lack of a consensus on what constitutes a “gold standard” for the representation, analysis, and interpretation of network data on ecological interactions within the framing of specific ecological questions; *i.e.*, which of the many available measures do actually hold ecological meaning? All things considered, the analysis of ecological networks can be confusing to newcomers as well as researchers who are more well-versed in existing methods.

Most notions in community ecology, including the definition of a community (Vellend 2010; Morin 2011), and several definitions of a niche (Holt 2009; Devictor et al. 2010), emphasize the need to study the identity of species and their interactions simultaneously (although ecological network analysis can be critiqued for ignoring species identity in many instances). Studies of ecological communities can therefore not discard or disregard interactions ({McCann} 2007), and using network theory allows researchers to achieve this goal. With the existence of methods that can analyze (large) collections of interactions, this approach is methodologically tractable. Graph theory (*e.g.* Dale & Fortin 2010) provides a robust and well formalized framework to handle and interpret interactions between arbitrarily large (or small) numbers of species. Theoretical analyses of small assemblages of interacting species (*e.g.* “community modules”, Holt 1997) have generated key insights on the dynamics of properties of ecological communities. We expect there is even more to gain by using graph theory to account for structure at increasingly high orders of organization (*e.g.* more species, larger spatial or temporal scales), because there is virtually no upper bound on the number of nodes (species) or edges (interactions) it can be applied to, and theory on large graphs can help predict the asymptotic behaviour of ecological systems. In short, although graph theory may appear as overwhelmingly complicated and unnecessarily mathematical, it allows us express a variety of measures of the structure of networks that can be mapped onto ecologically relevant questions.

Applying measures from network science to ecological communities can open three perspectives (Poisot et al. 2016d). First, the multiplicity of measures confers additional tools to *describe* ecological communities. This can reveal, for example, unanticipated ways in which communities differ. Second, these measures can provide new explanatory variables to *explain* how ecological communities function. The question of stability, for example, has been approached through the analysis of empirical food webs to question long-standing theoretical results (Jacquet et al. 2016). Finally, and this is a new frontier in network studies, they open the ability to *predict* the structure of ecological communities, through the prediction of interactions (Desjardins-Proulx et al. 2017; Stock et al. 2017). The domain of application of ecological networks is as vast as the domain of application of community ecology; but ensuring that network measures deliver their full potential of advancing our understanding of ecological systems require that they are well understood, and well used. Because of advances in graph theory, and the availability of more efficient computational methods, the exploration of large networks is now feasible. While this may not be immediately useful to macrobe-based research, microbial ecology, through sequencing, is able to generate datasets of immense size, that can be analyzed with the tools we present here (Faust & Skvoretz 2002).

This manuscript provides an assessment of the state of methodological development of network science applied to ecological communities. Taking stock of the tools available is necessary to determine how we can best analyze data from ecological networks. Previous work reviewed the consequences of network structure on ecological properties of communities and ecosystems (see Jordano & Bascompte 2013 for mutualistic systems, Poulin (2010) for parasites, {McCann} (2012) for food webs, or Dormann et al. (2017) for a recent overview), and we will not return to these topics. Instead, we highlight areas in which future research is needed, so as to eventually establish a comprehensive framework for how ecological networks can be analyzed. The measures presented in this manuscript do not represent all the measures that are available for ecological networks; instead, they represent a core set of measures that are robust, informative, and can be reasoned upon ecologically. While this manuscript in itself is not the entire framework for ecological network analysis, we are confident that it provides a solid foundation for its future development, and that the recommendations we lay out should be used by future studies. We have organized the measures by broad families of ecological questions. What is the overall structure of ecological networks? How can we compare them? What are the roles of species within networks? How similar are species on the basis of their interactions? How can we assess the significance of measured values? What are emerging questions for which we lack a robust methodology? This order mimics the way networks are usually analysed, starting from community-level structure, and going into the species-level details.

## 1. WHAT ARE SPECIES INTERACTION NETWORKS?

Identifying interactions across ecological entities can be done in a variety of ways, ranging from literature survey and expert knowledge, direct or indirect observation in the field using gut content (Carscallen et al. 2012), stable isotopes, molecular techniques such as meta-barcoding and environmental DNA (Evans et al. 2016; O’Donnell et al. 2017), to modelling based on partial data or mechanistic models. Depending on how they were assembled, species interaction networks can represent a multitude of ecological realities. When based on field collection (Morand et al. 2002; Bartomeus 2013; Carstensen et al. 2014), they represent *realized* interactions, known to have happened (unreported interactions can be true or false absences, depending on sampling effort among other things). Another common method is to “mine” the literature (*e.g.* Havens 1992; Strong & Leroux 2014) or databases (*e.g.* Poisot et al. 2016c), to replace or supplement field observations. In this situation, species interaction networks describe *potential* interactions: knowing that two species have been observed to interact once, *there is a chance* that they interact when they co-occur. Another more abstract situation is when interactions are inferred from a mixture of data and models, based on combinations of abundances (Canard et al. 2014), body size (Gravel et al. 2013; Pires et al. 2015), or other traits (Crea et al. 2015; Bartomeus et al. 2016). In this situation, species interaction networks are a *prediction* of what they could be. In keeping with the idea of “networks as predictions”, a new analytical framework (Poisot et al. 2016b) allows working directly on probabilistic species interaction networks to apply the family of measures presented hereafter.

Interactions are compiled and resolved (and subsequently assembled in networks) for a multitude of taxonomic and organisational levels (Thompson & Townsend 2000): individuals (Araújo et al. 2008; Dupont et al. 2009, 2014; Melián et al. 2014); species (Morand et al. 2002; Krasnov et al. 2004); at heterogeneous taxonomic resolutions, including species, genera, and more diffusely defined “functional” or “trophic” species (Martinez et al. 1999; Baiser et al. 2011); groups of species on the basis of their spatial distribution (Baskerville et al. 2011). This is because species interaction networks are amenable to the study of all types of ecological interactions, regardless of the resolution of underlying data: mutualistic, antagonistic, competitive, and so on. Recent developments made it possible to include more than one type of interaction within a single network (Fontaine et al. 2011; Kéfi et al. 2012), which allows greater ecological realism in representing communities, which encompass several types of interactions (*e.g.*, plants are consumed by herbivores, but also pollinated by insects). Such networks are instances of *multigraphs* (in which different types of interactions coexist). Another development accounts for the fact that ecological interactions may have effects on one another, as proposed by *e.g.* Golubski & Abrams (2011); these are *hypergraphs*. Hypergraphs are useful when interactions rely, not only on species, but also on other species interactions: for example, an opportunistic pathogen may not be able to infect a healthy host, but may do so if the host’s immune system is already being compromised by another infection. Hence it is not only species, but also their interactions, which interact. As using these concepts in ecological research represents a recent development, there is little methodology to describe systems represented as multigraphs or hypergraphs, and we will only mention them briefly going forward. In a way, methodological developments on these points is limited by the lack of data to explore their potential. As the interest among network ecologists will increase for systems in which the current paradigm of species–species interactions falls short, we expect that the inflow of data will stimulate the emergence of novel methods.

Formally, all of these structures can be represented with the formalism of graph theory. A graph G is defied as an ordered pair (V, E), where every element of E (the edges) is a two-element subset of V (the nodes). From this simple structure, we can measure a large number of properties (see e.g. Newman 2010 for an introduction). A simple graph contains neither self-edges (a node is linked to itself) or multiedges (the same two nodes are linked by more than one type of edge), whereas a *multigraph* contains at least one multiedge. As we illustrate in {fig. 1}, edges can be *directed* (*e.g.* A eats B), or *undirected* (*e.g.* A and B compete); *unweighted* (*e.g.* A pollinates B) or *weighted* (*e.g.* A contributes to 10% of B’s pollination). In the context of studying ecological interactions, is a set of ecological objects (taxonomic entities, or other relevant components of the environment), and *E* are the *pairwise* relationships between these objects. As both the strengths of interactions and their direction are highly relevant to ecological investigations, data on species interactions are most often represented as *networks*: directed and weighted graphs. We use network as a shorthand for “graph” throughout. Species interaction networks can, finally, be represented as *unipartite* or *bipartite* networks. *Unipartite* networks are the more general case, in which any two vertices can be connected; for example, food webs or social networks are unipartite (Post 2002; Dunne 2006). Unipartite networks can represent interactions between multiple groups; for example, food webs can be decomposed in trophic levels, or trophic guilds. Bipartite networks, on the other hand, have vertices that can be divided in disjointed sets T (*top*) and B (*bottom*), such that every edge goes from a vertex from T, to a vertex from B; any ecological community with two discrete groups of organisms can be represented as a bipartite network (parasites and hosts, Poulin 2010; e.g. plant and mutualists, Jordano & Bascompte 2013; phage and bacteria, Weitz et al. 2013). It is possible to represent k-partite networks, i.e. networks with k discrete ‘levels’. This formalism has been used for resources/consumers/predators (Chesson & Kuang 2008), and other plant-based communities (Fontaine et al. 2011). Tripartite networks are usually analyzed as collections of bipartite networks, or as unipartite networks.There still exists little data on ecological k-partite networks, and it is therefore dificult to establish solid recommendation about how they can be analyzed; this is a part of the field in which methodological development is still needed and ongoing.

**Figure 1:**
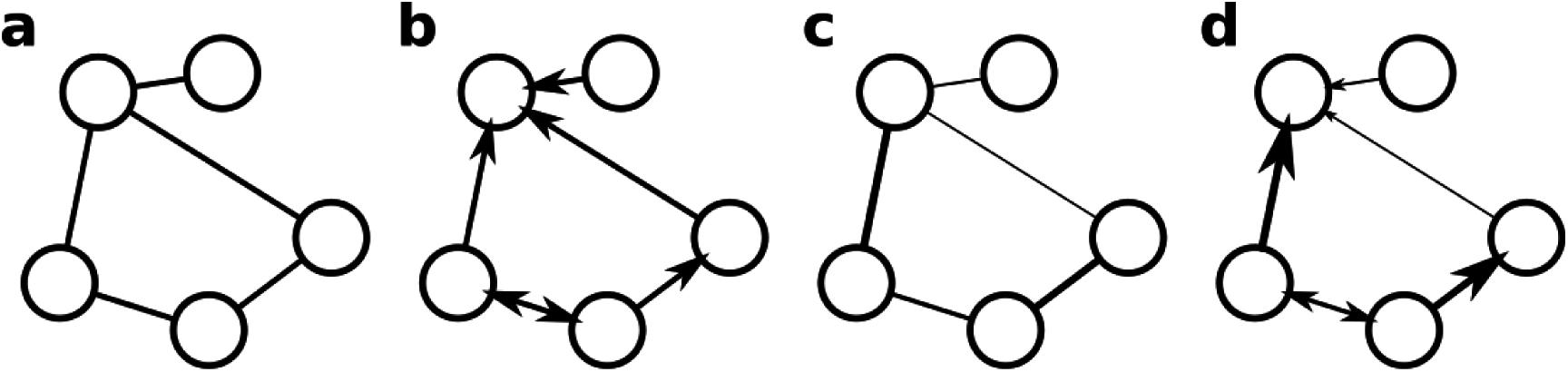
Differences between (un)weighted and (un)directed graphs. Graphs A and C are undirected, and graphs A and B are unweighted.

Networks can be represented using their *adjacency matrix* (**A**). For a unipartite network containing *S* species, **A** is a square matrix of dimensions (*S, S*). For a bipartite network containing T+B species, the dimensions are (*T, B*), and the **A** matrix is usually referred to as the *incidence* matrix. In both cases, the elements *a*_*ij*_ of the matrix indicate whether species *i* interact with species *j*. In unweighted networks, *a*_*ij*_ = 1 when *i* and *j* interact and 0 otherwise. In weighted networks the strength of the interaction is given, instead of being set to unity. Note that in weighted networks, the strength of the interaction is not necessarily between 0 and 1; if the strength of interactions depicts the raw effect of one population on another, then it can take on both negative and positive values. The adjacency matrix is symmetrical for undirected networks, because *a*_*ij*_ = *a*_*ji*_. In simple networks, the matrix diagonal is empty as there are no self-edges (which, ecologically, could represent autophagy, breastfeeding in mammals or cannibalism). We would like to note that A is not the *de facto* community matrix: in some situations, it can be more profitable to describe the community using its Jacobian matrix, *i.e.* one in which represents the net effect of species *i* on species *j* (Gravel et al. 2016b; Monteiro & Faria 2016; Novak et al. 2016), and therefore provides insights into the dynamics the system is expected to exhibit.

## 2. WHAT CAN WE LEARN WITH ECOLOGICAL NETWORKS?

For this part, unless otherwise stated, we will focus on describing measures of the structure of unweighted, directed networks (*i.e.* either the interaction exists, or it does not; and we know which direction it points), to the exclusion of quantitative measures that account for the *strength* of these interactions. In most of the cases, quantitative variations of the measures we present exist (see *e.g.* Bersier et al. 2002), and share a similar mathematical expression. We think that focusing on the simplifying (yet frequently used) unweighted versions allows one to develop a better understanding, or a better intuition, of what the measure can reveal. There is a long-standing dispute (Post 2002) among ecologists as to whether “arrows” in networks should represent biomass flow (*e.g.* from the prey to the predator) or interaction (*e.g.* from the predator to the prey). Because not all interactions involve biomass transfer, and because networks may be used to elucidate the nature of interactions, we will side with the latter convention. In general, we will assume that the interaction goes *from* the organism establishing it *to* the one receiving it (*e.g.* from the pollinator to the plant, from the parasite to the host, etc).

### 2.1 How do species interact in a community?

#### 2.1.1 Order, size and density

During the last decades, various network measures have been developed to characterize the general structure of interacting communities, capturing both species identity and their interactions (Dunne et al. 2002b; Montoya et al. 2006; Allesina & Pascual 2007; Thompson et al. 2012). Most of these measures encompass and supplement usual measurements in community ecology. In addition to how many species are there, and which species are in local area, knowledge of their interactions is indeed an additional layer of information that network measures exploit to quantify biodiversity.

A first descriptor of a network is its *order* (*S*), *i.e.* the total number of nodes. If nodes are species, order measures the species richness of the community described by the network *G*. The total number of inter<u>a</u>ctions (*L*) is the *size* of the network. From these two measures is computed the *linkage density*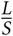 (*e.g.* Bartomeus 2013), which is the mean number of interactions per node – or simply, if a random species is selected, how many interactions would it be expected to have. Linkage density should be considered with caution as it can be misleading: the distribution of interactions among nodes in species interaction networks is rarely uniform or normal (Williams 2011), and a minority of species are known to establish a majority of interactions (Dunne et al. 2002a). Moreover *L* is known to scale with *S*^2^ (Cohen & Briand 1984; Martinez 1992), at least in trophic interaction networks.

This observation that *L* scales with *S*^2^ has cemented the use of an analog to linkage density, the *connectance* (*Co*), as a key descriptor of network structure (Martinez 1992). Connectance is defined as 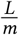, *i.e.* the proportion of established interactions (*L*), relative to the possible number of interactions *m*. The value of *m* depends of the type of network considered. In a unipartite directed network, *m* is *S*^*2*^. In a directed network in which species cannot interact with themselves, *m* is *S*(*S*-1). In an undirected network, *m* is 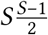 if the species cannot interact with themselves, and 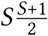 if they can. In a bipartite network, *m* is *T* × *B*, the product of the number of species at each level. The connectance varies between 0 if the adjacency matrix is empty to 1 if its entirely filled. It is also a good estimate of a community sensitivity to perturbation (Dunne et al. 2002a; Montoya et al. 2006) as well as being broadly related to many aspects of community dynamics (Vieira & Almeida-Neto 2015). Although simple, connectance contains important information regarding how links within a network are distributed, in that many network properties are known to strongly covary with connectance (Poisot & Gravel 2014; Chagnon 2015), and the fact that most ecological networks “look the same” may be explained by the fact that they tend to exhibit similar connectances ({fig. 2}). Poisot & Gravel (2014) derived the *minimum* number of interactions that a network can have in order for all species to have at least one interaction. This allows us to express connectance in the [0; 1] interval, where 0 indicates that the network has the least possible number of interactions.

**Figure 2.**
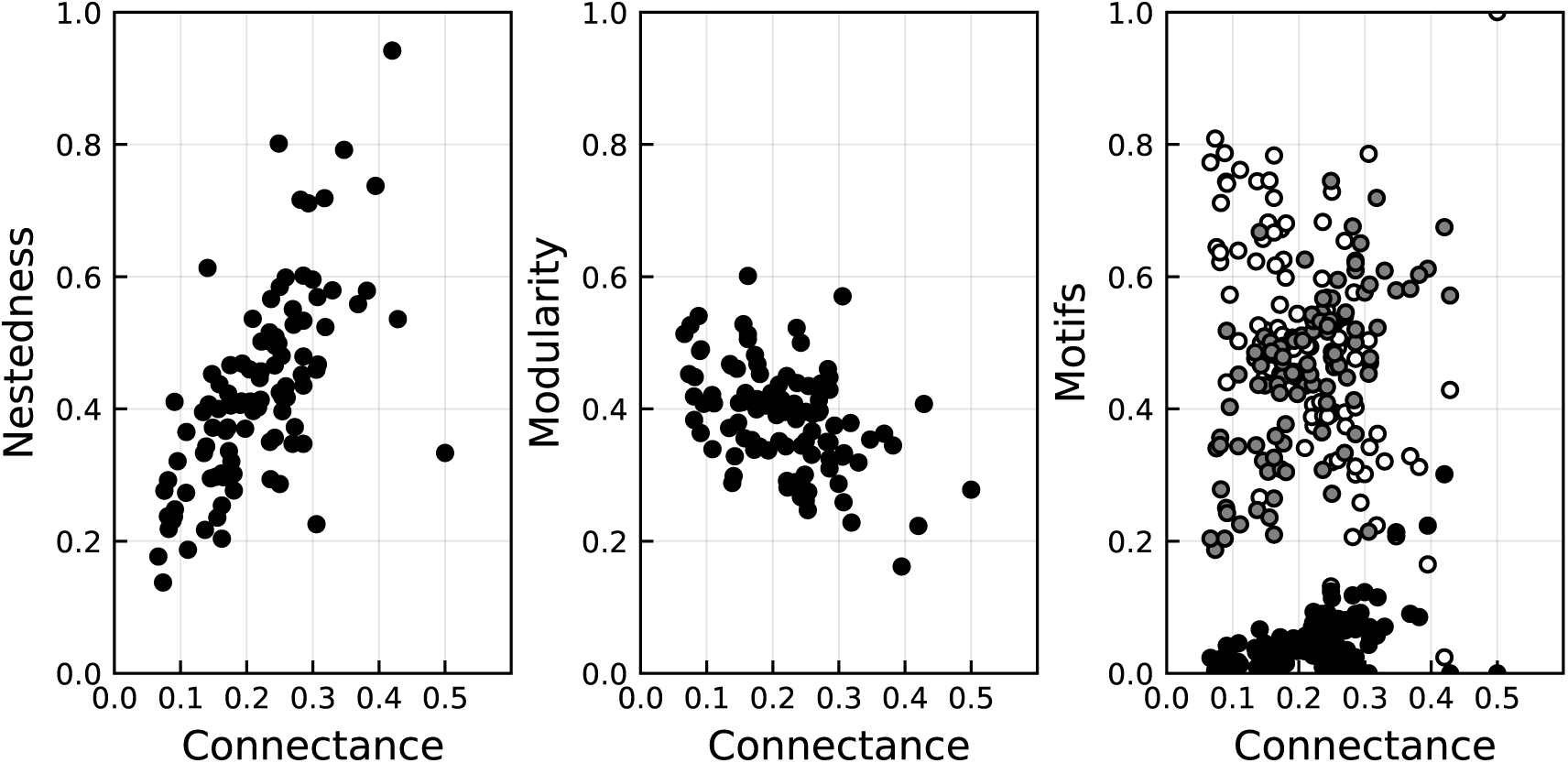
To illustrate the strong relationship betweeen connectance and other network measures, we measured the nestedness using η, modularity (best partition out of 100 runs), and the relative frequencies of three bipartite motifs (sparsely connected, white; partially connected, greyl fully connected, black) in 102 pollination networks. The sparsely connected motif represents two independant interactions. The partially connected motif represents the addition of one interactions to the sparsely connected one, and the fully connected is the addition of another interaction. All of these measures have a strong covariance with connectance, and for this reason, the comparison of networks with different connectances must rely on randomizations. Data, methods, and code: **https://osf.io/82ypq/**.

#### 2.1.2 Interactions repartition within the networks

The majority of real-world species interaction networks are highly heterogeneous with regard to interactions distribution among nodes. This distribution can be studied as such (through the *degree distribution*), but also reflects a particular organization of the network, which can also be studied. Quantitative measures of different structures have been developed from graph theory and have played a growing role in understanding the evolution and functioning of ecological communities – in particular, because these measures add a small amount of information (comparatively to measures presented later in this manuscript), they are a natural first step in moving away from a species-centric view of community into the arguably more realistic species-and-interactions view that networks capture well.

The *degree* of a node is its number of interactions, then the *degree distribution P* (*k*) measures the probability that a species has *k* interactions within the network. The degree distribution can be calculated as *P* (*k*) = *N* (*k*)/*S* where *N* (*k*) is the number of nodes with *k* interactions, and S is the number total of species in the network. The degree distribution allows identification of important nodes, such as potential keystone species (Solé & Montoya 2001; Dunne et al. 2002b), generalists, and specialist species (Memmott et al. 2004). In directed networks, the degree distribution can be divided into *in-degree* and *out-degree*. These respectively correspond to species *vulnerability* (*e.g.* number of predators in food webs) and *generality* (*e.g.* number of resources in food webs). It is often assumed that the distribution of degree in networks should resemble a power law (Strogatz 2001; Caldarelli 2007). In other words, the proportion P(*k*) of nodes with degree *k* should be proportional to *k*^−γ^ (but see see Jordano et al. 2003 – a truncated power-law may be a more accurate description). Assuming that power laws are an appropriate benchmark is equivalent to assuming that ecological networks are structured first and foremost by preferential attachment, and that deviation from power law predictions suggests the action of other factor. Dunne et al. (2002a) found that, at least in food webs, ecological networks tend not to be small-world or scale-free (*i.e.* having a specific degree distribution; Caldarelli 2007), but deviate from these rules in small yet informative ways (specifically, about prey selection or predator avoidance). Opportunistic attachment and topological plasticity have been suggested as mechanisms that can move the system away from predictions based on power laws s (Ramos-Jiliberto et al. 2012; Ponisio et al. 2017). We suggest that deviations from the power law be analysed as having intrinsic ecological meaning: why there are more, or fewer, species with a given frequency of interactions may reveal reasons for and/or constraints on particular species interactions.

The network *diameter* gives an idea of how quickly perturbations may spread by providing a measure of how dense the network is. Diameter is measured as the longest of all the shortest *distances* (*d*_*ij*_) between every pairs of nodes in the graph (Albert & Barabási 2002), where *d*_*ij*_ is the length of the shortest path (sequence of interactions) existing between the nodes *i* and *j*. A small diameter indicates presence of a densely connected nodes, or hubs, hence fast propagation between nodes which may make the network more sensitive to perturbation (*e.g.* rapid spread of a disease, Minor et al. 2008). The diameter is relative to the number of nodes in the network, since it relies on counting the number of interactions in a path, which may become larger as the network order increases. To overcome this issue, the diameter can also be measured as average of the distances between each pair of nodes in the network.

#### 2.1.3 Aggregation of nodes based on their interactions

From the heterogeneous repartition of interactions between nodes in species interaction networks, certain structures and grouping of interactions around nodes emerge. While the degree distribution hints at how interactions are organized around single nodes, one can frame this question at the scale of the entire network. It is likely that other structure will appear when multiple nodes are considered at once. This can be done by analyzing what types of relationships the nodes (representing species, etc) are typically embedded in (*e.g.* competition, intraguild predation), through the analysis of *motifs distribution*, or by determining if there are nodes found in dense *clusters* or non-overlapping *compartments*, forming *modular* communities.

Species interaction networks can be decomposed in smaller subgraphs of species, called motifs (Milo et al. 2002). The smallest modules to which they can be decomposed are three-species motifs (Holt 1997). The relative frequencies of each of these motifs holds information about network structure. There are thirteen possible three-nodes motifs in directed networks, each representing a different relationship between three nodes, such as competition between A and B for a shared resource C (*A* → *C* ← *B*), or a linear chain between A, B and C (*A* → *B* → *C*). Among these thirteen motifs, some are present in species interaction networks with a lower or higher frequency that what is expected in random networks. Motifs distributions are characteristic of network type (neuronal, electrical, social, ecological, and so on). In food webs for example, motifs’ under- and over-representation has been found to be consistent across different habitats (Camacho et al. 2007; Stouffer et al. 2007; Borrelli 2015). In ecological networks, motifs have been referred to as the *basic building blocks of communities*, as they represent typical relationship between species. Studying their distribution (*i.e.* how many of each type of motif is there is this network) offers an opportunity to bridge the gap between two traditional approaches (Bascompte & Melián 2005), namely the study of the dynamics of simple modules such as omnivory or linear food chain (Pimm & Lawton 1978; Holt 1996; {McCann} et al. 1998), and the analysis of aggregated metrics describing the community as a whole. Motif distributions have been used to study the processes underlying the assembly and disassembly of ecological communities (Bastolla et al. 2009), as well as of the link between communities’ structure and dynamics (Stouffer & Bascompte 2011). More recently, motifs have also been used to define species trophic roles in the context of their community (Baker et al. 2014) and link this role to the network’s stability (Borrelli 2015).

The *clustering coefficient* is useful to estimate the “cliquishness” of nodes in a graph (Watts & Strogatz 1998) – that is their grouping in closely connected subsets. It measures the degree to which the neighbours of a node are connected (the neighborhood of a node *i* is composed of all of the nodes that are directly connected to *i*). In other words, it gives an idea of how likely it is that two connected nodes are part of a larger highly connected group or “clique”. Two different versions of the clustering coefficient (*CC*) exist. First, it can be defined locally, for each node *i (*Watts & Strogatz 1998). In this case 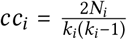 where *k*_*i*_ is *i*’s degree (its number of neighbors) and *N*_i_ is the total number of interactions between *i*’s neighbors. It describes the fraction of realized interactions between *i*’s neighbors and thus vary between 0 (none of *i*’s neighbors are connected) and 1 (all of them are connected, forming a “clique”). From this measure, we can calculate the average local clustering coefficient: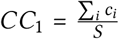 where is the total number of nodes. This first version describes the “cliquishness” of a typical neighborhood, but has the drawback of giving more influence to nodes with a small degree. Nevertheless, the clustering coefficient provides a way of characterising the structure of the graph through the analysis of *CC _k_*, which is the average of the *cc*_*i*_ of all nodes of degree *k*, and specifically of the distribution of *CC*_*k*_ across multiple values of *k*. The clustering coefficient can also be defined globally, for the entire graph (Soffer & Vazquez 2005; Saramäki et al. 2007) and is calculated as follows 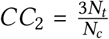, where *N*_*t*_ is the number of triangles in graph *G* (*a* is connected to *b* and *c, b* to *a* and *c* and *c* to *a* and *b*) and *N*_*c*_ is the number of 3-nodes subgraphs (*e.g. a* is connected to *b* and *c*, *b* and *c* are connected to *a* but not to each other). Kim (1993) suggested that this property of a network can be used to infer competition, but this has to our knowledge received little attention in ecology.

Whereas clustering analysis gives information about the grouping of nodes within their immediate neighbourhood (but no information about the identity of nodes in this neighborhood), a measure of modularity gives a similar information at a larger scale (Gauzens et al. 2015). Network modularity measure how closely connected nodes are divided in *modules*, also called *compartments* (Olesen et al. 2007). A module is defined as a subsystem of non-overlapping and strongly interacting species. The modular structure of graphs has been studied because of its dynamical implications, in that modularity promotes stability by containing perturbations within a module, thereby constraining their spreading to the rest of the community (Stouffer & Bascompte 2010, 2011). This has been a key argument in the diversity-stability debate (Krause et al. 2003). A major challenge when studying species interaction networks’s modularity is to find the best subdivision of the network. Several methods have been developed for this purpose, including the optimization of a modularity function (Guimerà et al. 2004; Newman 2004; Newman & Girvan 2004; Guimerà & Amaral 2005; Guimerà & Nunes Amaral 2005). The principle underlying this function is to find the optimal subdivision that maximizes the number of interactions within modules while minimizing the number of interactions between modules. The calculated modularity is then compared with a null model that has the same number of links and nodes, with the links connected to each other randomly. Modularity optimization has a resolution limit (in that its performance decreases with the size of the network) making it less reliable for large species interaction networks (Fortunato & Barthelemy 2007); there are methods designed specifically to work on thousands of nodes and more (see *e.g.* Liu & Murata 2009). To compare outcomes of different modularity measurements, it possible to use an *a posteriori* method. In a network where modules are already found, the realized modularity (*Q*´*R*) measure the proportion of interactions connecting nodes within modules (Poisot 2013). This is expressed as

**Figure 3.**
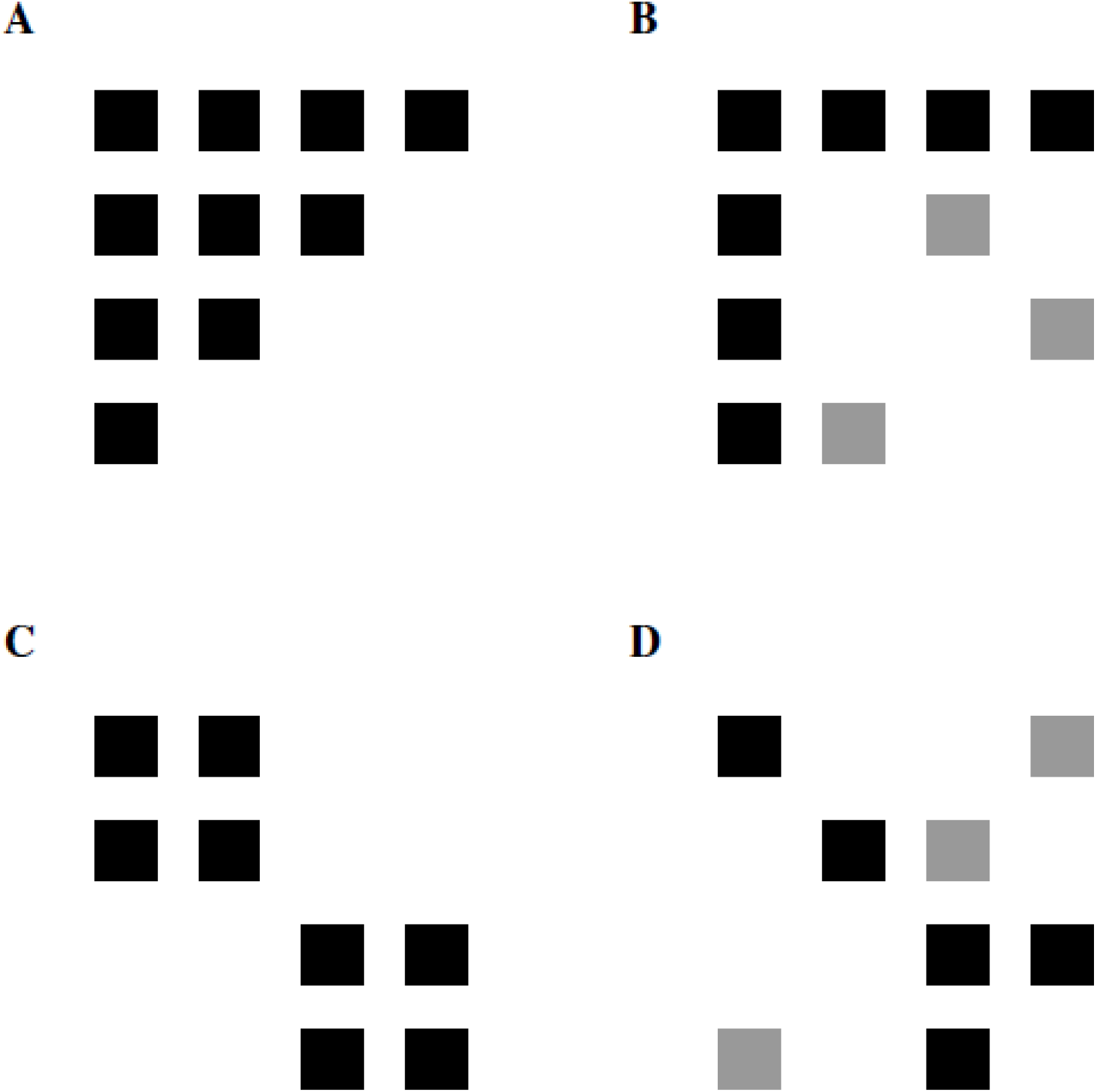
Illustration of the nested and modular structure of networks, represented as matrices. A is a perfectly nested matrix; in B, three interactions (in grey) have been displaced to lose the perfectly nested structure. C is a perfectly modular network; in D, three interactions have been displaced to lose the modular structure.

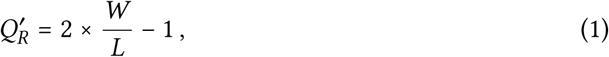

where *W* is the number of interactions *within* modules, and *L* is the total number of interactions. This takes on a value of 1 when modules are disconnected from one another (which is not true of other modularity functions that account for the probability of establishing an interaction). This measure can take on negative values if there are more interactions between modules than within them, which can be viewed as a non-relevant partitioning of the community.

#### 2.4.1 Nestedness

Species interaction networks can also present a *nested* structure, where the species composition of small assemblages are subsets of larger assemblages. In food webs, a nested structure occurs when the diet of the specialists species is a subset of the diet of the more generalist species – and where the predators of species are nested as well. The analysis of nestedness has revealed ecological and evolutionary constrains on communities. For example, it has been hypothesized that a nested structure promotes a greater diversity by minimizing competition among species in a community (Bastolla et al. 2009). Various metrics have been developed to quantify nestedness (Ulrich 2009; Ulrich et al. 2009). Most are based on the principle that when a matrix is ordered by rows and columns (that is descending in rank from above and from the left) a nested networks will present a concentration of presence data in the top-left corner of the matrix, and a concentration of absence data in the opposite corner (see Staniczenko et al. 2013 for an exception). Numerous studies (Rodriguez-Girones & Santamaria 2006; Fortuna et al. 2010; Flores et al. 2011) use the proportion of unexpected presence or absence in the matrix to quantify nestedness. Seemingly the most widely used measure of nestedness is NODF (nestedness measure based on overlap and decreasing fills), as suggested by Almeida-Neto et al. (2007); Bastolla et al. (2009) suggested that η can complement NODF, in that η does not require a re-ordering of the nodes (*i.e.* there is no need to put the most densely connected nodes first, and the least densely connected nodes last). As per Bastolla et al. (2009), η is defined as:

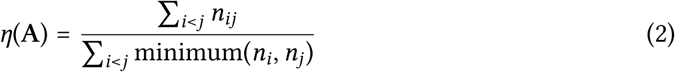

where *n*_*ij*_ is the number of common interactions between species *i* and *j*, and *n*_*i*_ is the number of interactions of species *i*. Note that this formula gives the nestedness of rows with regard to the columns, though one can also measure the nestedness of columns with regard to rows as η (A’), and calculate the nestedness of the whole system as the average of these two values. We suggest that, since it does not rely on species re-ordering, η can be used over NODF or other nestedness measures. There are some caveats to this argument, however. First, the number of permutations for NODF is known, and for species-poor networks, they can be computed in a reasonable time. Second, NODF can help understanding how different orderings of the matrix (*e.g.* using traits instead of degree) contributes to nestedness – if this is the question of interest then NODF is the logical choice (Krishna et al. 2008). Once ordered by degree, NODF and η are identical (with the exception that NODF accounts for decreasing fill, whereas η does not).Finally, η has the undesirable property of always giving the same value depending only on the degree distribution. Therefore, any permutation of a network that maintain the degree distribution will give the same value of η, which greatly impedes hypothesis testing.

#### 2.1.5 Intervality

A last measure of species interaction networks structure is their intervality. The first step in calculating intervality is to identify a common trait along which nodes can be ordered. This can be body mass in the case of food webs, but can also be a property derived from their position in the network, such as their degree; indeed, a nested bipartite network is interval when species are organized by decreasing degree. Intervality then measures how well interactions of all species can be described by this trait. A network is called *interval* when it can be fully explained by one dimension (trait). An interval food web with species ordered by their body mass, as an example, has predator eating a consecutive range of preys, that all fall into a range of body masses (Eklöf & Stouffer 2016), or are closely related from a phylogenetic standpoint (Eklöf & Stouffer 2016). Most unipartite ecological networks are close to being interval with one or several dimensions, such as defined by body size (Zook et al. 2011) or arbitrary traits derived from the interactions themselves (Eklöf et al. 2013). There are several methods to quantify a network’s intervality. Cattin et al. (2004) have measured the “level of diet discontinuity” using two measures: (i) the proportion of triplet (three species matrix) with a discontinuous diet (*i.e.* at least one species gap), in the whole food web (*D _diet_*), and (ii) the number of chordless cycles 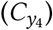. A cycle of four species is considered as chordless if at least two species out of the four are not sharing prey, so the diets cannot be totally interval. Nevertheless, these two measures only give a local estimation of the intervality. Stouffer et al. (2006) proposed to measure the intervality of the entire network by re-organizing the interaction matrix to find the best arrangement with the fewer gaps in the network. This is a stochastic approach that by definition does not guarantee to find the global optimum, but has the benefit of working at the *network* scale rather than at the scale of triplets of species.

### 2.2 How are communities different?

Detecting spatial and temporal variation in ecological networks, and associating these variations to environmental factors, may yield insights into the underlying changes in ecosystem functions, emergent properties, and robustness to extinction and invasion (Tylianakis et al. 2007; Tylianakis & Binzer 2013). These efforts have been hindered by the difficulty of quantifying variation among interaction networks. The challenge lies in finding a meaningful way to measure the dissimilarity between networks (Dale & Fortin 2010). Given the ecological properties or processes of interest, a direct comparison – not always computationally tractable – may not be necessary. Hence, networks can be indirectly compared through their properties (*e.g.* degree distribution, connectance, nestedness, modularity, etc.). Multivariate analyses of network metrics have been used to estimate the level of similarity between different networks (Vermaat et al. 2009; Baiser et al. 2011), while null models were used to statistically compare observed values to their expected random counterparts (*e.g.* Flores et al. 2011).

In the situation where several networks share a large enough number of species, one can alternatively compare how these shared species interact. This approach can be particularly useful alongside environmental gradients (Tylianakis et al. 2007; Tylianakis & Morris 2017). It represents a second “dimension” of network variability, where in addition to changes in higher order structure, changes at the scale of species pairs *within* the networks are accounted for. This variation is more readily measured through a different approach to sampling, where instead of relying on the sampling of a large number of networks in different environments, efforts are focused on the same system at reduced spatial or temporal scales. The development of methods to analyse replicated networks is still hampered by the lack of such data; this is especially true in food webs. Replicated food webs based only on the knowledge of the local species and their potential interactions (*e.g.* Havens 1992) are not always appropriate: by assuming that interactions always happen everywhere, we do not capture all sources of community variation (in addition to the issue of co-occurrence being increasingly unlikely when the number of species increases). Sampling of ecological networks should focus on the replicated documentation of interactions within the same species pool, and their variation in time and space (Poisot et al. 2012; Carstensen et al. 2014; Olito & Fox 2015), as opposed to relying on proxies such as comparison of different communities across space (Dalsgaard et al. 2013), or time (Roopnarine & Angielczyk 2012; Yeakel et al. 2014).

Analysis of network structure measures has so far played a central role in the comparison of networks and in the search for general rules underpinning their organization (Dunne 2006; Fortuna et al. 2010). Notably, the number of species affects the number of interactions in real ecological networks (Martinez 1992; Brose et al. 2004), and thus many other network properties (Dunne 2006). Some measures of network structure covary with expected ecological properties, such as species abundance distributions (Blüthgen et al. 2008; Vázquez et al. 2012; Canard et al. 2014), network size and sampling intensity (Martinez et al. 1999; Banašek-Richter et al. 2004; Chacoff et al. 2012). This issue can seriously limit the interpretation of network measures and their use for network comparison. Furthermore, most of these measures are highly correlated among themselves: Vermaat et al. (2009) report that network variation can be reduced largely along three major axes related to connectance, species richness (which is tied to connectance because the number of interactions scales with the number of species) and primary productivity (which is hard to measure, and is not easily defined for all systems). More recently, Poisot & Gravel (2014) and Chagnon (2015) showed that because of constraints introduced by the interaction between connectance and network size, the covariation of the simplest measures of network structure is expected to be very strong. As a consequence, it is barely possible to make robust network comparisons using the variations in these basic descriptors. We therefore need to go beyond these global network properties, and find meaningful alternatives that allow a better understanding of the ecological differences between networks.

#### 2.2.1 Higher order differences in structure

Other methods accounting for the structure of the entire network have been developed. For example, some methods are based on the frequency distribution of small subnetworks including network motifs (Milo et al. 2002) and graphlets (a more general definition of motifs; Przulj 2007; Yavero lu et al. 2015). The method of graph edit distance gives edit costs (each modification to the graph counts for one unit of distance) for relabeling nodes, as well as insertion and deletion of both nodes and interactions (Sanfeliu & Fu 1983), and therefore provides a well-defined way of measuring the similarity of two networks (this method has not been widely used in ecology). Other suitable measures to determine network similarity are based on graph spectra (Wilson & Zhu 2008; Stumpf et al. 2012). Spectral graph theory (which is yet to be applied comprehensively to the study of species interaction networks, but see Lemos-Costa et al. (2016)) characterizes the structural properties of graphs using the eigenvectors and eigenvalues of the adjacency matrix or the closely related Laplacian matrix (the Laplacian matrix, defined as **D – A**, wherein **D** is a matrix filled with ‘s in the off-diagonal elements, and the degree of each node on the diagonal, accounts both for network structure and for degree distribution). Some methods allow the algorithmic comparison of multiple networks in which no species are found in common (Faust & Skvoretz 2002; Dale & Fortin 2010), and are primarily concerned about the overall statistical, as opposed to ecological, properties of networks.

#### 2.2.2 Ecological similarity and pairwise differences

All of the aforementioned methods focus on the mathematical similarity of networks rather than their ecological similarity. To fill this gap, Poisot et al. (2012) presented a framework for measurement of pairwise network dissimilarity, accounting both for species and interactions turnover through space, time or along environmental gradients. Following Koleff et al. (2003), this approach partitions interactions in three sets: shared by both networks, unique to network 1, and unique to network 2. The β-diversity can be measured by comparing the cardinality of these three sets to reflect symmetry of change, gain/loss measures, nestedness of interaction turnover, etc. This method of network. β-diversity can also be extended to multiple network comparisons using their relative difference from the same meta-network. While many measures of β-diversity exist to analyse compositional data, there is still a lack of a comprehensive methodology regarding their applications to networks. A large part of the issue stems from the fact that species interactions require the species pair to be shared by both communities, and consequently some analyses require that the species pair is shared by two communities: measures of network β-diversity are strongly constrained by the structure of species co-occurrence. In no species pairs co-occur, or if no two networks have common species, these methods cannot give informative results (the dissimilarity being, by default, complete) – as of now, this suggests that a tighter integration of these methods with research on compositional turnover is needed, especially to understand the threshold of shared species below which they should not be applied. In addition, none of the current methods seem sufficient for characterizing the structure for a meaningful comparison and extracting information hidden in the topology of networks (as they ignore network-level structure, *i.e.* emerging from more than direct interactions), and the development of future methods that work regardless of species composition seems like a straightforward high-priority topic. Finally, this framework would benefit from a better integration with quantitative measures. Using Bray-Curtis (or equivalent) measures to assess difference between networks for which interaction strengths are known would allow to quantify dissimilarity beyond the presence or absence of interactions.

### 2.3 What do species do?

Not all species in large communities fulfill the same ecological role, or are equally important for processes and properties acting in these communities. As species interactions are a backbone for fundamental mechanisms such as transfer of information and biomass, one can expect that the role of a species reflects its position within its community, organized by trophic level, abundance, body size or other ecologically meaningful organizing principle. In species interaction networks, it is possible to measure the position and the role of species in different ways, giving different ecological information.

#### 2.3.1 Centrality

Centrality is a measure of how “influential” a species is, under various definitions of “influence”. It has been used to identify possible keystone species in ecological networks (Jordán & Scheuring 2004; Martín González et al. 2010). We would like to note that the ability of network structure measures to identify keystone species is highly dubious; the canonical definition of a keystone species (Paine 1969) requires knowledge about biomass and effects of removal, which are often not available for network data, and make predictions that are primarily about species occurrences. These measures may be able to identify list of candidate keystone species, but this requires careful experimental / observational validation. Nevertheless, knowledge of network structure allows us to partition out the effect of every species in the network. For example, in networks with a nested structure, the core of generalist species have higher centrality scores, and the nested structure is thought to play an important role for network functioning and robustness (Bascompte et al. 2003). We provide an illustration of some centrality measures in {fig. 4}. In this section, we review five measures of centrality: degree, closeness, betweenness, eigenvector, and Katz’s.

**Figure 4.**
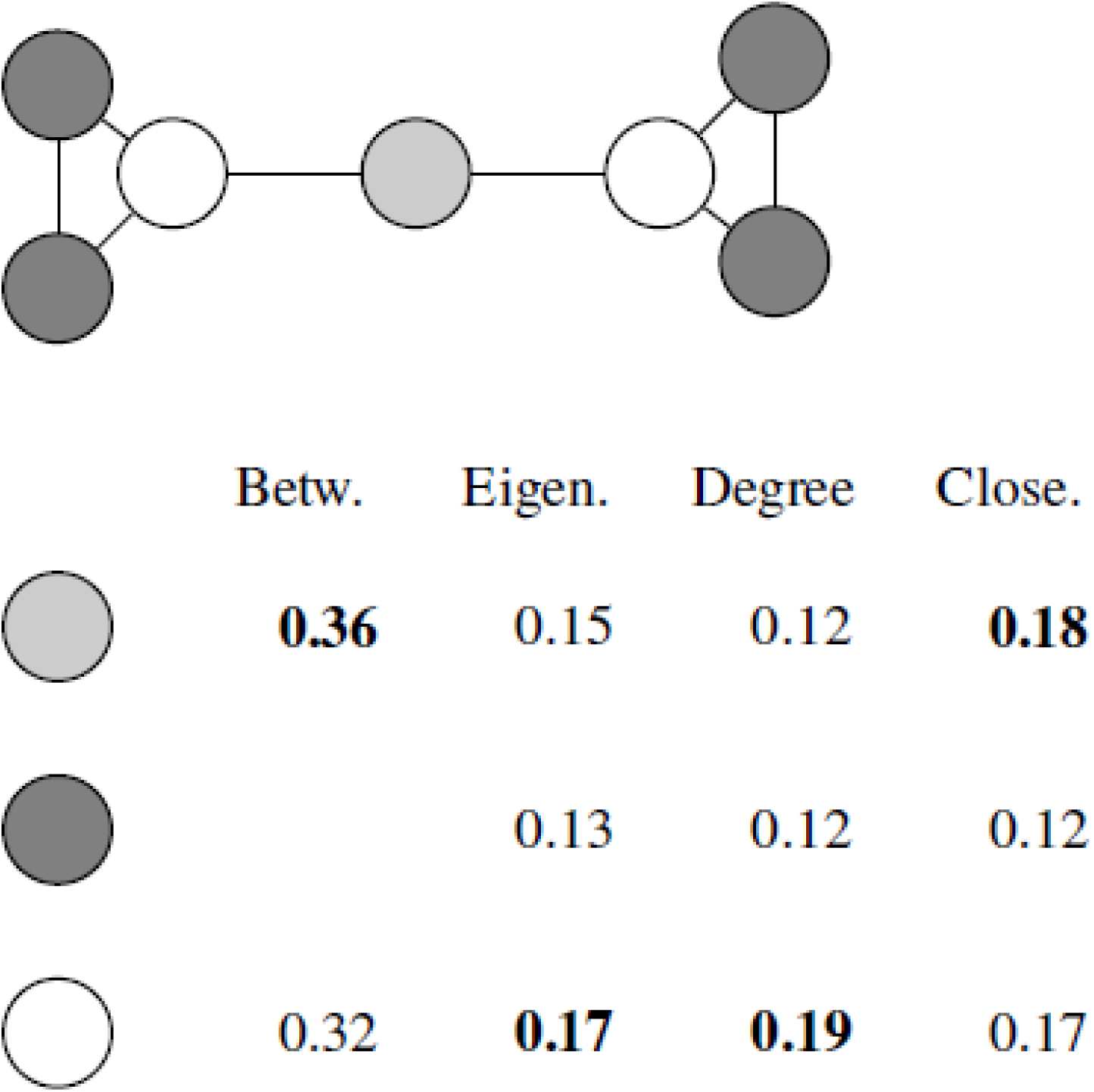
On the simple graph depicted in the top (nodes of the same color have the same centralities), we measured centrality using betweenness, eigen centrality, degree centrality, and closeness. The values have been corrected to sum to unity. The value in bold gives the most central family of nodes for the given measure. This example illustrates that different measures make different assumptions about what being “central” mean. The dark grey nodes do not have a betweenness centrality value; some software will return 0 for this situation.

Degree centrality (*C*_*D*_(*i*) = *k*_*i*_; Freeman (1977)) is a simple count of the number of interactions established by a species. In directed networks, this measure can be partitioned between in-degree (interactions from others to *i*) and out-degree (interaction from *i* to other). It is a *local* measure, that quantifies the immediate influence between nodes. As an example, in the case of a disease, a node with more interactions will be more likely both to be infected and to contaminate more individuals (Bell et al. 1999). To compare species’ centrality, *C*_*D*_ has to be normalized by the maximum degree (〈*C*_*D*_〉 = *C*_*D*_/*k*_max_).

Closeness centrality (*C*_*C*_) (Freeman 1978; Freeman et al. 1979) measures the proximity of a species to *all* other species in the network, and is therefore *global* in that, although defined at the species level, it accounts for the structure of the entire network. It is based on the shortest path length between pairs of species and thus indicates how rapidly/efficiently a node is likely to influence the overall network. The node with the highest *C*_*C*_ is closer to all other node than any other nodes and will thus affect more rapidly the overall network if, for example, there is a perturbation (Estrada & Bodin 2008). Formally, *C*_*C*_ is defined as

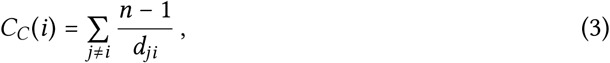

where *d*_*ij*_ is the shortest path length between *i* and *j*, and *n* is the number of species.

Betweenness Centrality (*C*_*B*_) (Freeman 1977) describes the number of times a species is *between* a pair of other species, *i.e.* how many paths (either directed or not) go through it. This measure is thus ideal to study the influence of species loss on fragmentation processes for example (Earn 2000; Chadès et al. 2011; McDonald-Madden et al. 2016). Nodes with high *C*_*B*_ values are considered as modules connectors in modular networks. The value of *C*_*B*_ is usually normalized by the number of pairs of species in the network excluding the species under focus, and is measured as

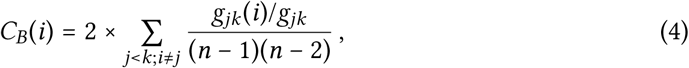

where *g _jk_* is the number of paths between *j* and *k*, while *g*_*jk*_(*i*) is the number of these paths that include *i*.

Eigenvector centrality (*C*_*E*_ – Bonacich 1987) is akin to a simulation of flow across interactions, in which each species influences all of its partners simultaneously. It then measures the relative importance of species by assigning them a score on the basis that an interaction with more influential species contribute more to a species’ score than the same interaction with a low-scoring species (Allesina & Pascual 2009). From a graph adjacency matrix A, the eigenvector centrality of species *i* is given by

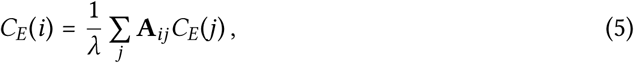

where **A** _*ij*_ is 1 if *i* interacts with *j* and 0 otherwise, and λ is a constant. This can be rewritten as the eigenvector equation:

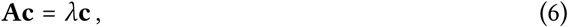

wherein **c** is the vector of all values of *C*_*E*_ As all values of *C*_*E*_ have to be positive, as per the Perron-Frobenius theorem, λ is the greatest eigenvalue of **A**.

Finally, Katz’s centrality (*C*_*K*_ – Katz 1953) is a measure of the influence of a node in the network. This measure takes into account all the interactions connecting a node to its neighborhood. However, an immediate neighbor has more weight than a distant one. *C*_*K*_ is defined as

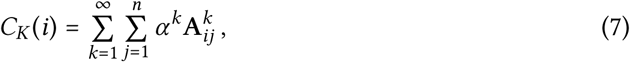

wherein α is the *attenuation constant*, and *k* is the length of the paths between *i* and *j*. The α value is between 0 and 1/λ, where λ is the largest eigenvalue of **A**. Larger values of α give more importance to distant connections, thus allowing this measure to function either locally (immediate neighborhood) or globally (entire graph). *C*_*K*_ can be used in directed acyclic graphs (*e.g.* trees), which is not true of *C*_*E*_. This is also the only measure to have a probabilistic equivalent (Poisot et al. 2016b).

Studying different measures of centrality provide important information regarding the roles of certain species/nodes. As an example, a species may have a low *C*_*D*_ and a high *C*_*B*_, meaning that it plays a key role in connecting species that would not be connected otherwise even if it does not interact with them directly. A low *C*_*D*_ and a high *C*_*C*_ means that the species has a key role by interacting with important species. Because the absolute values of centrality vary with network size and connectance, Freeman et al. (1979) suggest that the *centralization* measure, rarely applied in ecology, be used when comparing centrality across networks. Centralization is defined, for any centrality measure *C*_*x*_, as the sum of the differences between each node’s centrality, and the highest centrality value (Σ_*i*_ (*C*_*x*_(*i*)-max(*C*_*x*_))). This measure is then divided by the maximal possible value of centralization for a network with the same number of nodes and interactions, which in turns depends on the formulae used to measure centrality, and can be estimated based on random draws of the networks.

#### 2.3.2 Species roles in the network

Species functional roles can be reflected in the interactions they establish (Coux et al. 2016), providing a clear bridge between network approaches and functional ecology studies. Functional traits are known to be correlated with the position of species in the network, either because they intervene directly in the interaction (Brose et al. 2006a; Alexander et al. 2013), constraining the set of possible interactions or their frequency, or because phenological incompatibilities prevent the interaction from happening (Olesen et al. 2011). For instance, Petchey et al. (2008a) used allometric scaling of body size and foraging behaviour of individual consumers to predict species interaction. Scaling up to multiple traits, one can group species into functional clusters, based on their similarity. The distribution of some species-level network measures (*e.g.* centrality, degree) can then be compared within and across groups (Petchey & Gaston 2002). This method usually does not account directly for interactions between species (Petchey et al. 2008a) but is useful when studying a process for which the influential traits are known, or to test the importance of a particular (complex of) traits on a function. Note that one *can*, in this situation, adopt a very generous definition of what constitutes a trait: spatial grouping of species (Baskerville et al. 2011) for example, is one example in which examining interactions in the light of species *attributes* provides ecological insights.

If external information on species traits is absent, the role of a species can be approached through the interactions it establishes within the network: species with similar interactions are often grouped in *trophic species*, and these can be assumed to have similar traits or lifestyles (this approach has mostly been used in food webs). Indeed, many food web models (Williams & Martinez 2000; Cattin et al. 2004) predict interactions between *trophic groups*, and not between species. Lumping species within trophic groups maintains the heterogeneity of interactions *across* groups, but removes all variability of interactions between species *within* the groups. As a consequence, species that bring unique interactions to a trophic group may be overlooked. Dalla Riva & Stouffer (2015) suggested an alternative to this approach: species positions are analyzed *before* clustering them into groups (*i.e.* there is a measure of position for every species), allowing explicit investigation of species interactions *while* avoiding obfuscation of the variance within groups.

Coux et al. (2016) measured the functional role of species, by applying (“functional dispersion”; Laliberté & Legendre 2010) to the adjacency or incidence matrix of the network. Under this framework, like in Mouillot et al. (2013), the uniqueness of a species is hinted at by its distance to the centroid of all other species. We argue that this approach should be questioned for two reasons. First, it is sensitive to the ordination choices made. Second, it is not clear how it allows the comparison of results across different networks: not only does the position of a species vary in relation to other species in the network, it varies from one network to another. Note that centrality measures are not necessarily better at identifying which species are unique: as we show in {fig. 4}, for some measures, non-unique nodes have high centrality values. We argue that the development of measures for node uniqueness should receive increased attention. In particular, measures that rely on ordination only account for first-order interactions, *i.e.* the direct interactions between species. As a consequence, a large part of the network structure, which emerges through consideration of longer chains of interactions, is not accessible via these methods.

Looking at network motifs is a promising way to address species functional roles and node uniqueness. Motifs are all the possible ways a fixed number of species (usually three or four) can interact. Within these motifs, species can occupy a variety of unique positions; for example, within a linear food chain, there are three distinct positions (bottom, middle, top), whereas a trophic loop has a single unique position. Within motifs with three species, 30 unique positions can be identified (Stouffer et al. 2012), and for each species, its frequency of appearance at each of these position within networks has been shown to be an inherent characteristic conserved through its evolutionary history. This method has the advantage of grouping species that may be different in terms of guild or partners, but that contribute in the same way to the structure of the community. Based on this vector it is possible to statistically identify species that exhibit similar profiles. Motif positions tend to be well conserved both in time (Stouffer et al. 2012) and space (Baker et al. 2014), making them ideal candidates to be investigated alongside functional traits and phylogenetic history.

#### 2.3.3 Partition based on modularity

In large communities, some species are organized in modules (see sec. 2.1.2), within which they interact more frequently among themselves than with species of the same overall network but outside of their module. Guimerà & Nunes Amaral (2005) proposed that when functional or topological modules can be found in large networks, the functional role of a species can be defined by how its interactions are distributed within its module and with other modules. To identify these roles, the first step is to identify the functional modules of a large network. The profile of species interactions is determined by using two measures.

First, the *z*-score quantifies how well-connected a species *i* is within its module *m*.

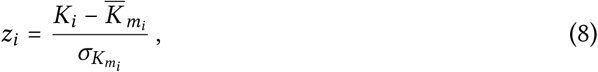

where *K*_*i*_ is the degree of *i* within its module *m*_*i*_; 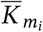is the average of K over all species of *m*_*i*_ and 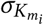 is the standard deviation of *K* in *m*_*i*_.

Second, the *participation coefficient* (PC) describes the profile of *i*’s interaction with species found outside of the module *m*.

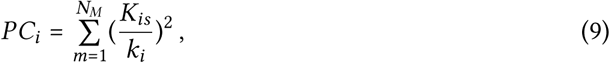

where *k*_*i*_ is the total degree of species *i*, meaning a count of all its connection, inter- and intra module. The *P C* of a species therefore varies between 0 (all interactions are within the module) and 1 (all interactions are uniformly distributed among all the modules). The use of these indices is based on the assumption that species with similar interactions have similar traits and so are expected to play the same functional role.

Olesen et al. (2007) use these two values to divide species in four groups, based on a cutoff for *z* (2.5) and for *P C* (0.62). Species with low *z* and low *P C* are “peripherals” – they are not well connected within or between modules. Species with low *z* and high *P C* connect well between, but not within, modules, and are “connectors”. Species with high *z* and low *P C* are “module hubs”, well connected within their own modules but not with the outside. Finally, species with high *z* and high *P C* are “network hubs”, connecting the entire community. In their analysis of plants and pollinators, Olesen et al. (2007) reveal that pollinators tend not to be module hubs, and are also less frequently network hubs than plants are.

#### 2.3.4 Contribution to network properties

As species make differential contributions to network structure and processes, the removal of certain species will therefore have a greater effect on the community’s stability and functioning, and these species are therefore stronger contributors to these processes. Differential contribution to several processes can be estimated in multiple ways: by performing removal/addition experiments in real ecological systems (*e.g.* Cedar creek or BIODEPTH experiments), by analyzing the effect of a species extinction within empirical (Estrada & Bodin 2008) or simulated (Berlow et al. 2009) systems, by using a modeling approach and simulating extinctions (Memmott et al. 2007), or by analyzing the statistical correlation between an ecosystem property and species functional roles (Thompson et al. 2012). Another way to quantify the contribution of a species to a property *P* is to compare it to its contribution to the same property when its interactions are randomized (Bastolla et al. 2009). This method allows studying the contribution of a species’ interactions, as the variation of interactions is intuitively expected to be faster than the variation of species. Indeed, because interactions require species to co-occur, because there are far more interactions than species, and because interactions have a dynamic of their own, whether there will be more signal in interactions than in species presence is an hypothesis that should be tested on empirical systems in priority.

The contribution of a species to a given network measure after its interactions are randomized is

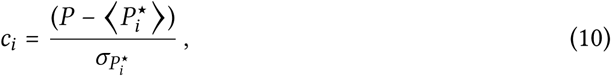

where *P* is the property (nestedness, modularity, productivity …), 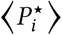and 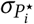are the average and standard deviation of the property across a set of random replicates for which species *i* interactions have been randomized. The effects of several traits or structural properties of species (such as centrality or STR) on their contributions to a given measure can then be analyzed.

### 2.4 How similar are species interactions?

Some species exhibit a much larger set of interactions than others or form denser clusters within the network. One of the many challenges of ecology is to understand the causes and consequences of such heterogeneous species interactions. Species are, first and foremost, related by their phylogenetic history. We will not address this aspect here, because it does not easily integrate with network theory. We encourage readers to refer to Cadotte & Davies (2016) instead.

One way in which the heterogeneity of species interactions is quantified is through analysis of the overlap in their partners, known as ecological similarity. For simplicity, we will use the vocabulary derived from trophic networks, but these methods can be applied to other types of ecological networks. Ecological similarity between species is a widely used concept that quantifies the resemblance between two species or “biotic interaction milieu” (McGill et al. 2006) and allows analyzing processes ranging from species niche (Elton 1927) and community assembly (Piechnik et al. 2008; Morlon et al. 2014) to trophic diversity (Petchey & Gaston 2002). The simplest and most widely used measure of pairwise ecological similarity is the Jaccard coefficient (Legendre & Legendre 2012):

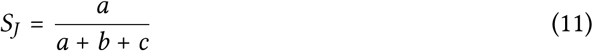

where *a* is the number of shared partners, *b* the number of species that interact with only the first species and *c* with only the second species (for variations, see (Legendre & Legendre 2012)). The Jaccard similarity coefficient is widely used to estimate ecological similarity and competition between species (Rezende et al. 2009) but does not account for the shared absence of interactions (but see Chao et al. 2005). This is not a severe issue, as ecological networks tend to be extremely sparse, and therefore shared absence of interactions may not be informative. The similarity index has to be chosen with care depending on the focus of the study. In the general equation above, consumers and resources are seen as perfectly equivalent (additively), but, in directed networks, it can be adapted to include consumer and resources as different dimensions of trophic activities and/or for dynamical food webs by including information about flows (Yodzis & Innes 1992). Once a similarity matrix is formed from all pairwise measurements, a hierarchical clustering can be performed to build a dendrogram, which gives information about the trophic diversity of species within a community and the relative uniqueness of species (but see Petchey et al. 2008b). Cophenetic correlation (Sokal & Rohlf 1962) can then be used to analyze how well several dendrograms, built using different methods, preserve the similarity between species (Yodzis & Winemiller 1999). The similarity of overall communities can also be estimated to see how similar, or dissimilar, species within it are when compared to null models (Morlon et al. 2014). For this purpose, the mean or maximum pairwise similarity is averaged across the whole network under consideration.

### 2.5 Is any of this significant?

Most network properties tend to be colinear, specifically because they covary with connectance. For example, the number of interactions in a network with a known number of species will limit the possible values of nestedness, modularity, and so on (Poisot & Gravel 2014). As such, the value of any measure of network structure often needs to be compared to a range of possible values under a null model. The purpose of the null model is to search the null space of possible randomized networks (Fortuna et al. 2010), in a way that would yield an unbiased distribution of the measure of interest, to which the observed value is then compared. In practice, this approach is constrained by (i) the size of the null space to search, and specifically the fact that it varies with connectance (Poisot & Gravel 2014), and (ii) the computational burden of a thorough null space exploration.

A large number of studies use the null hypothesis significance testing (NHST) paradigm to assess the significance of an observed value of network structure. NHST works by generating *randomized* networks under a variety of constraints, measuring the property of interest on these randomizations, then commonly using a one-sample *t*-test with the value of the empirical measure as its reference. This is justified because, through the mean value theorem, the application of enough randomizations should yield a normal distribution of the simulated network measure (see Flores et al. 2011). Bascompte et al. (2003) used a probabilistic sampling approach, where the probability of drawing an interaction depends on the relative degree of the species; Fortuna & Bascompte (2006) used the same approach, with the distinction that all interactions have the same probability (equal to connectance). Drawing from a probability distribution in this manner has a number of shortcomings, notably the fact that some species can end up having no interactions, thus changing the network size (which Fortuna et al. (2010) termed “degenerate matrices”). An alternate approach is to use constrained permutations, where pairs of interactions are swapped to keep some quantity (the overall number of interactions, the degree of all species, and so on) constant. This approach is used in null models for species occupancy (Gotelli 2000; Gotelli & Entsminger 2003; Ulrich & Gotelli 2007). Stouffer et al. (2007) used an intermediate approach, where swapping was done as part of a “simulated annealing routine”, to give the algorithm enough leeway to explore non-optimal solutions before converging (as opposed to just swapping, which has no definition of the optimality of a solution). As of now, there are no clear recommendations as to which approach to sample the null space is the most efficient (or computationally feasible for large network sets), emphasizing the need for a more exhaustive comparison of the behaviour of these methods.

#### 2.5.1 Hypotheses underpinning topological null models

The most frequently used null models are *topological, i.e.* they can search the null spaced based only on the matrix, and do not rely on ecological processes to generate random networks. We will focus on the subset of null models which generate a probability of observing an interaction based on different aspects of network structure; these probabilistic networks can be analyzed directly, or as is most commonly done, converted into binary networks through random draws. There are three broad categories of null models (commonly used for bipartite networks) – based on connectance, based on degree distribution, and based on marginal degree distribution. Each family embodies a specific hypothesis about the sources of bias on the measured property.

Type I (Fortuna & Bascompte 2006) null models are focused on *connectance*, where the probability of any two species *i* and *j* interacting is fixed as

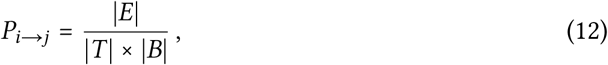

where *T* and *B* are vertices from the “top” (*T* = { υ ∈ *V, k*_in_(υ) = 0}) and “bottom” (*B* = { υ ∈ *V,k_out_* (υ)=0}) levels of the network (these methods where originally applied to bipartite networks). This model assumes that interactions are distributed at random between all species, without considering the degree of the species. Deviation from the predictions of this model indicate that the network measure of interest cannot be predicted by connectance alone.

Type II (Bascompte et al. 2003) null models add an additional level of constraint, in that it respects the degree distribution of the network (in degree *k*_in_; out-degree *k*_out_)In a Type II network,

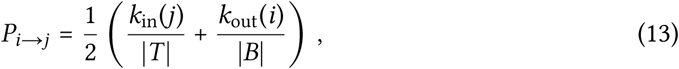

meaning that the interaction is assigned under the hypothesis that *i* distributes its outgoing interactions at random, and *j* receives its incoming interactions at random as well. In this model, species with more interactions have a higher probability of receiving interactions in the simulated network. This respects both the distribution of generality and vulnerability. Deviation from the predictions of this model indicate that the network measure of interest cannot be predicted by the degree distribution alone.

Finally, Type III null models account for only one side of the degree distribution, and can be defined as Type III in, wherein

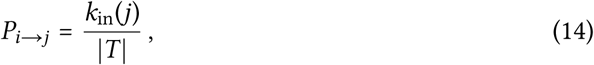

and Type III out, wherein

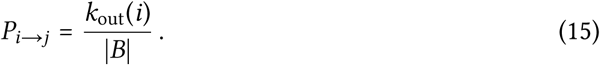

Deviation from the predictions of this model indicate that the network measure of interest cannot be predicted by the marginal degree distributions alone. Ecologically speaking, deviation from this null model means that the way interactions are established / received is sufficient to explain the observed structure. These models can be expressed in a sort of hierarchy. Type I introduces the least hypotheses, and should be applied first. If there is no significant deviation, then Type III models can be applied, then Type II. This approach has the important benefit of, in addition to determining which properties show a difference from the random expectation, give insights about which aspect of the structure are responsible for this difference.

#### 2.5.2 Topological and generative models

It is important to note that these models, based on permutations, are purely *topological*. There is no difference, when deciding if an interaction should be assigned between two species, between *e.g.* a plant-pollinator network, or a host-parasite network. One may want to test deviation from a null distribution that would be informed by ecological processes. To inject some processes in the null models used, several “generative” models have been proposed. By contrast to topological models, generative models use core assumptions about ecological mechanisms to generate networks that mimic aspects of a template network. Arguably the most influential (despite it being limited to trophic interactions) is the “niche model” (Williams & Martinez 2000), that generates networks of trophic groups based on the hypothesis that feeding interactions are determined by an arbitrary niche-forming axis generally accepted or implied to be body-size ratios (Brose et al. 2006a). Gravel et al. (2013) showed that the parameters of this model can be derived from empirical observations. The niche model assumes a beta distribution of fundamental niche breadth in the entire network (in cases where the trait space in bound between 0 and 1); this assumption, close though it may be to empirical data, nevertheless has no mechanistic or theoretical support behind it. This suggests that so-called generative models may or may not be adequately grounded in ecological mechanisms, which implies the need for additional developments. Similar models include the cascade model and the nested-hierarchy model, but these tend to generate networks that are qualitatively similar to those of the niche model (Brose et al. 2006b). More recently, several models suggested that species traits can be used to approximate the structure of networks (Santamarίa & Rodrίguez-Gironés 2007; Bartomeus 2013; Crea et al. 2015; Olito & Fox 2015; Bartomeus et al. 2016). Finally, networks tend to be well described only by the structure of species abundances. Both in food webs (Canard et al. 2012) and host-parasite bipartite networks (Canard et al. 2014), modelling the probability of an interaction as the product of relative abundance is sufficient to generate realistic networks. These generative models represent an invaluable tool, in that they allow building on mechanisms (though, as we illustrate with the niche model, not necessarily ecological ones) instead of observed relationships to generate the random expectations. The NHST-based analyses then proceeds as with topological models, *i.e.* the observed value is compared to the distribution of values in the theoretical space.

### 2.6 Future methods for novel questions

Surveying the methodological toolkit available to analyze ecological networks highlights areas in which future developments are needed. We identified, in particular, four topics that would require additional attention.

### 2.6.1 Multi/hyper graphs

Most of the tools to analyse species interaction networks are limited to node-to-node interactions, to the exclusion of node-to-interaction or interaction-to-interaction interactions. This limits the variety of biological situations that can be represented. Golubski & Abrams (2011) presented a number of situations that elude description in this way. For example, opportunistic infection by a pathogen *O* requires the pre-existence of an interaction between a pathogen *P* and an host *H*. This situation is better captured as (i) the existence of an interaction between *H* and *P*(noted *L*_*HP*_) and (ii) the existence of an interaction between *O* and this interaction, noted *O* → *L*_*H P*_. Another hard-to-represent scenario is niche pre-emption: if a host can be infected by either pathogen *P*_1_ or *P*_2_, but not both at the same time, then the interactions 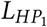and 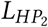interact antagonistically. This is a different situation from simple competition between *P*_1_ and *P*_2_. Although these are extremely important drivers of, for example, species distributions (Araújo & Rozenfeld 2014; Blois et al. 2014), the current methodological framework of ecological network analysis is not well prepared to represent these data.

### 2.6.2 External information

Building on the basis suggested by Poisot et al. (2015), Bartomeus et al. (2016) proposed that the mechanisms determining ecological interactions can be identified within a cohesive statistical framework, regardless of the type of ecological interaction. At its core, their framework assumes that interactions are the consequence of *matching rules, i.e.* relationships between traits values and distributions. For example, a pollinator can get access to nectar if its proboscis is of a length compatible with the depth of the flower. Rather than relying on natural history, these “linkage rules” (Bartomeus 2013) can be uncovered statistically, by modelling an interaction L_*ij*_ as a function *f* (*x*_*i*_, *y*_*j*_) of the traits involved, wherein *x*_*i*_ and *y*_*j*_ are sets of traits for species *i* and *j* respectively. Procedures akin to variable selection will identify the traits involved in the interaction, and model selection can identify the shape of the relationship between trait values and interactions. There are two reasons for which this work is an important milestone in the modern analysis of ecological networks. First, it places interactions within the context of community ecology, by showing how they build upon, and influence, trait distributions. In particular, it draws attention to how structure of networks results both from the linkage rules and from the distribution of traits in the locality where the network is measured (Gravel et al. 2016a). Second, it does away with the necessity of topological models to generate random networks: identifying matching rules is the only step needed to generate random networks based on *functional, biological* hypotheses, thereby solving some of the concerns we identified with generative null models. We argue that this approach should be expanded to accommodate, *e.g.* phylogenetic relationships between species (Bastazini et al. 2017). The ideal framework to study networks, and the one we should strive for, avoids considering interactions in isolation from other aspects of community structure – instead, it is explicit about the fact that none of these aspects are independent. Although this will come with additional mathematical and statistical complexity, this cost will be more than offset by the quality and the refinement of the predictions we will make.

Although documenting species, traits, and interactions seems like a daunting effort, there are novel approaches to accelerate the generation of data in some systems. For example, Bahlai & Landis (2016) show that passive measurement based on citizen science (using Google Images) allows users to accurately document phenological matches and species interactions between flowers and bumblebees. Similarly, Evans et al. (2016) show that sequencing of diet gives access to phylogenetic and interaction history within a single experiment. Addressing novel questions will require a diversification of the methodological toolkit of network ecologists, as well as an improved dialog between empiricists and theoreticians.

### 2.6.3 Networks of networks

An additional frontier for methodological development has to do with the fact that networks can be nested. A network of species–species interactions is the addition of interactions at the population level (Poisot et al. 2015), themselves being aggregates of interactions at the individual level (Dupont et al. 2011, 2014; Melián et al. 2014). This is also true when moving from single-site to multi-site network analysis (Poisot et al. 2012; Canard et al. 2014; Carstensen et al. 2014; Trøjelsgaard et al. 2015). Local interaction networks exist in meta-community landscape (Gravel et al. 2011; Trøjelsgaard & Olesen 2016), and their structure both locally and globally, is constrained by, but is also a constraint on, co-occurrence (Araújo et al. 2011; Cazelles et al. 2015).

Analyzing networks in a meta-community context might require a new representation. Most of the challenge comes from two sources. First, species are shared across locations; this means that two nodes in two networks may actually represent the same species. Second, networks are connected by species movement. Both the dynamics and the structure of networks are impacted by the fact that species move across the landscape at different rates and in different ways. The implication is that every species in the landscape potentially experiences its own version of the metacommunity (Olesen et al. 2010). These issues have seldom been addressed, but would allow a more potent examination of the spatial structure and dynamics of ecological networks (Trøjelsgaard & Olesen 2016). Gravel et al. (2016b) recently introduced spatially explicit Jacobian matrices, allowing the formal consideration of coupled dynamics of several networks in a meta-community.

## 3. WHAT ARE SPECIES INTERACTION NETWORKS, REVISITED?

The above analyses benefit from access to (context-enhanced) data on ecological interactions. An important point to raise is that the format expected for the analysis (*i.e.* when data are actively being processed) is different from the format suitable for storage, archival, mining, and linking. From an information management perspective, this puts the question of *What are ecological networks?* in a new light.

Most of the measures mentioned above, and therefore most software, expect networks to be represented as matrices; every row/column of the matrix is an object, and the value at row *i* and column *j* is a measure of the interaction between *i* and *j*. It can be a Boolean value, a measure of interaction strength, or a probability of interaction. This approach is used by databases such as IWDB, Web-of-Life.es, and World of Webs (Thompson et al. 2012). Although this approach has the benefit of being immediately useful, it lacks the easy addition of metadata. In the context of species interaction networks, metadata is *required* at several levels: nodes (species, individuals), interactions, but also the overall network itself (date of collection, site environmental data, and so on). Most research has so far been *constrained* to the adjacency matrix representation of networks. However, ontologically richer representations (graphs with built-in metadata) may offer themselves to a larger and different tool set: multi-graphs, and hyper-graphs, capture a wider picture of ecosystems where all types of interactions are considered simultaneously. Food webs, or other networks, stored as binary or weighted matrices may not be the most relevant representation for these questions.

There are two initiatives that remedy this shortcoming by providing meta-data-rich information on ecological interactions. globi (Poelen et al. 2014) is a database of interactions, extracted from the literature, and available through *GBIF*. It relies on an ontology of interaction types, and on unique taxonomic identifiers for species. mangal.io (Poisot et al. 2016a) is a database of networks, that can be fed and queried openly through several packages; it relies on a custom data format, and can be linked to other databases through the use of taxonomic identifiers.

Networks formatted as raw matrices may well be immediately usable, but supplementing them with external information is hard. On the other hand, granular databases with rich metadata can always be converted to raw matrices, while retaining additional information. It is important that we maintain a distinction between the formats used for *storage* (in which case, relational databases are the clear winner) from the formats used for *analysis* (that can be generated from queries of databases). In order to facilitate synthesis, and draw on existing data sources, it seems important that the practice of depositing interaction matrices be retired, in the profit of contributing to the growth of context-rich databases. There are a handful of software packages available for ecological network analysis (Csardi & Nepusz 2006; Dormann et al. 2008; Hagberg et al. 2008; Hudson et al. 2013; Flores et al. 2016; Poisot et al. 2016b). They differ in their language of implementation, license, and methods availability.

Considerations about the analysis of networks go hand in hand with the far more difficult question of data sources and data quality. Jordano (2016) shows that obtaining estimates of the completeness of sampling is both difficult, and different between weighted and unweighted networks. Describing the data at the level of the *interaction* in more detail may therefore give better estimates of (i) the robustness of the overall network, and (ii) the relevant aspects of life-history to add in models. These can then be added to predictive models, in the form of functional traits (Bartomeus 2013; Bartomeus et al. 2016), to boost our ability to infer the existence of interactions (or their strength). Relevant interaction-level data (discussed in Poisot et al. 2016a) include the identity of species involved, their abundances, local environmental conditions, and functional traits of the individuals or populations observed interacting, when available. Shifting the focus of sampling away from networks, and onto interactions (because what are networks, but a collection of interactions?) would give more information to work with. Because the amount, resolution, and type of information that it is necessary and feasible to sample will vary for each system, empirical network scientists should lead the effort of developing data standards. Taking a step back, *data quality* should be framed within the context of a specific analysis; we feel that there is a need for a review that would attempt to determine the minimal amount of information needed as a function of the type of analyses that will be applied.

## 4. CONCLUSIONS

In this contribution, we have attempted a summary of the measures from graph theory that are the most frequently used in, or the most relevant for, the analysis of species interaction networks. Even though species interaction networks are ubiquitous in community ecology, biogeography, macroecology, and so on, there is no clear consensus on how to analyse them. We identified a number of areas that would benefit from more methodological development. We highlight each of them below, and identify whether they should stimulate future development of novel methods to complete the framework, or stimulate further investigation and assessment of existing methods to clarify when they should be applied.

1. There is a pressing need to accommodate hypergraphs and multigraphs within the SIN analysis framework, so as to work on a larger and more realistic variety of ecological situations. Pilosof et al. (2017) identified these systems as having a high relevance when predicting community change, and the emergence of zoonotic diseases, and this is a clear example of an area in which ecology and applied mathematics can have a fruitful interaction.

2. The information we use in the building of SINs needs to be expanded. Far from being a collection of species and their interactions, networks are structured by environmental forces, species traits distribution, species evolutionary history, and random chance. Replicated datasets with extensive metadata and additional information would most likely boost our power to describe, explain, and predict network structure (Poisot et al. 2016d). The next generation of network measures should account for additional information carried by both species and interactions.

3. Of course, the addition of data to ecological interactions requires to expand the scope of what is currently being sampled, and to normalize it to some extent. More broadly, we expect that the development of novel methods, and the collection of novel data and their standardization, should go hand in hand. The emergence of interactions and networks databases, based around documented formats, is a step in the right direction, as they provide an idea of the scope of data to collect.

4. We need to establish stronger standards for the manipulation of network data. Networks are difficult to manipulate, and the lack of a robust software suite to analyse them is a very worrying trend – our knowledge of ecological networks is only as good as our implementation of the analyses, and academic code can always be made more robust, especially in fields where the widespread adoptions of computational approaches is still ongoing. We expect that, since there are numerous initiatives to increase good practices in software engineering among academics, this problem will be solved by improved community standards in the coming years.

5. The NHST approach to network structure needs additional study, especially when it comes to determining best practices. Recent developments in graph theory, and notably edge-sampling based cross-validation (Li et al. 2016), can help assess the performance of generative null models. There is a shortage of null models that are based on topology but still account for known biology of the networks (such as forbidden interactions), highlighting the need for future developments.

6. There is a need to compare the alternative measures of a single property. We tried as much as possible to frame these measures in the context of their ecological meaning. But this can only be properly done by strengthening the ties between network analysis and field or lab based community ecology. Statistical analysis of measures on existing datasets will only go so far, and we call for the next generation of studies aiming to understand the properties of network structure to be built around collaboration between empirical researchers and measures developers.

## Acknowledgements

ED, MB, and TP contributed equally to this manuscript. Order of authors MB and ED was decided by flipping a coin. All authors besides MB, ED, MHB, GVDR and TP listed in alphabetic order. MB, ED, MHB, and TP wrote the article outline. ED and MB wrote the first draft. ED, MB, and TP revised the manuscript. All other authors contributed to edits and discussion. This work was conducted as a part of the Ecological Network Dynamics Working Group at the National Institute for Mathematical and Biological Synthesis, sponsored by the National Science Foundation through NSF Award #DBI-1300426, with additional support from The University of Tennessee, Knoxville.

